# Co-occurrence network multicollinearity artefacts using centered log-ratio transformation

**DOI:** 10.64898/2026.10.01.756072

**Authors:** Dennis D. Krutkin, Scott T. Kelley

## Abstract

Next-generation sequencing data is compositional in nature and ignoring this fact can significantly impact downstream data analyses. This is especially true in co-occurrence (correlation) networks because compositional data is prone to many false negative correlations. The centered log-ration (clr) transformation is a commonly used compositional data analysis (CoDA) approach that converts compositionally constrained data into real number space. The clr transformed values can subsequently be used in machine learning and statistical analyses such as Spearman or Pearson correlations used to generate co-occurrence networks. Here we show that the necessary zero substitution performed prior to clr transformation can create numerous multicollinearity artefacts. We also show that a newer CoDA approach called the robust centered log-ratio (rclr) eliminates these artefacts in correlation-based network analysis.

---

There is increasing recognition that next-generation sequencing data used for amplicon library sequencing, metagenomics and RNA sequencing, as well as other types of data (e.g., total ion normalized metabolomics data) are compositional in nature (1). Ignoring the compositionality of these data can significantly affect data analysis and downstream interpretation. In compositional data, the feature values within samples (e.g., proportions or sequence counts) sum to a fixed total meaning that an increase in the amount of one feature necessarily results in a reduction of all the other features (1, 2). Compositional data, therefore, violate the assumption of variable independence, a fundamental assumption of most statistical analyses.

In the 1980s, Atchison solved the issue of compositionality by proving that log-ratio transformation converts compositionally constrained data into real number space (3). This work led to the development of compositional data analysis (CoDA) methods first used in geology (rocks are compositions of mineral) and later expanded to other data (2). One of the most used and flexible transformations developed by Atchison is the centered log-ratio (clr) transformation in which the value of every feature in a sample is transformed into a natural log-ratio of the geometric mean of all the features in the given sample (4). The clr transform is simple to calculate and exceptionally useful for combining multiomics data on different scales, such as metagenomic and metabolomics data (2, 5, 6). Numerous studies have shown that such transformations, collectively termed compositional data analysis (CoDA), improve the ability to discriminate sample types and reduce artefacts due to compositionality (7–9).

One of the challenges with log-ratio transformations is the issue of zero values since the log of zero is undefined. This issue is especially problematic in large highly sparse data sets with many zero values that are common in multiomics datasets (10–13). Researchers often remove the sparse features from their datasets to reduce their impact (e.g., by removing features present in <10% of the samples). However, this still leaves many missing features. Zero-handling in clr transformation typically involves replacing zero values with a small, fixed value (e.g., 1 or 0.01) prior to transformation (2). These non-zero values are also clr transformed.

An alternative to the clr, called the robust clr (rclr) transform takes a different approach by ignoring zero values when performing the transformation (9). The rclr transform calculates the geometric mean of all non-zero values within a sample, and then only transforms the non-zero values by taking their log-ratio relative to the geometric mean. Post-transformation, former zero values can then be replaced with a fixed value (e.g., the minimum value of the entire dataset) (14).

The effects of clr and rclr transformation have primarily been tested for their effects on multidimensional scaling (9). Less attention has been paid to their usefulness in the context of correlation network analysis. The utility of CoDA analysis with correlation networks is clear, given the inherent problem of false negative correlations in compositional data (6). Many previous studies have pointed out the issues of spurious correlations with compositional data and have developed methods to overcome this problem when generating co-occurrence networks (15–17) including the use of clr transformation prior to performing Pearson or Spearman correlation (18–22).

Here we show that, while the clr transformation solves one source of network correlation artefacts (false negative correlations), zero-replacement prior to transformation can create novel spurious correlations (Fig. 1). Since every sample in a dataset has a unique set of features values, the geometric mean is different for each sample. This means that even if the zero-replacement values were the same for the entire dataset prior to transformation, they will all be slightly different after clr transformation (Fig. 1A). A closer look at the data underlying these networks show that these high correlations involve highly sparse features (mostly zero values in the original data set). Ironically, the sparser the data, the greater the likelihood of artefactual multicollinearity correlation between features. In rclr transformation any zero replacement occurs after log-ratio transformation which means that all the formerly zero values will be the same post rclr transformation (Fig. 1B).

**Figure 1.**
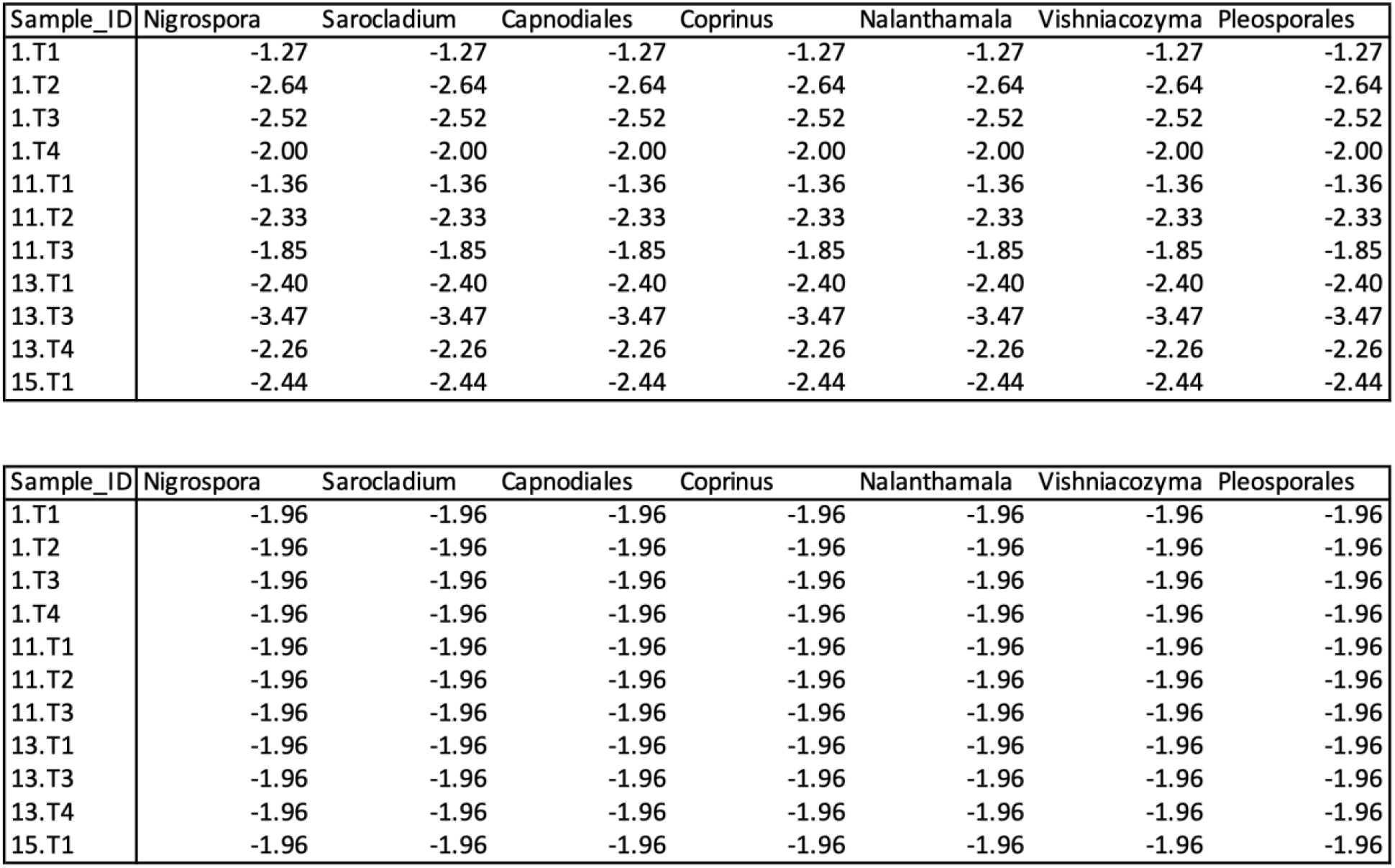
Example post-transformation values from the same subset of fungal ITS feature count data from Nannini et al. (24) for zero-replacement followed by clr transformation (TOP) and rclr transformation followed by minimum value replacement (BOTTOM). The transformations are based on the same dataset and of 7 features all of which had zero count values for in the 11 samples shown. In the clr transformed data, the zeros had been replaced by 0.01 prior to transformation and each sample ended up with a different value replacement post transformation. In the rclr transformed data, all the values were the same after minimum value replacement post-transformation.

While this does not appear to have a very strong impact on multidimensional scaling, it can create significant multicollinearity artefacts in correlation networks. Figure 2 compares of two correlation (co-occurrence) networks generated from the same dataset in figure 1 using either clr and rclr transformation. The network based on clr transformation includes a dense region of highly correlated edges that are not present in the rclr transformed network. This is occurring because in the clr transformation, the transformed zero values in one sample correlate with transformed zero values in other samples, which is completely nonsensical. Since all the values among samples post clr transformation are all the same within a sample but different among samples, this results in highly correlated features within the samples.

**Figure 2.**
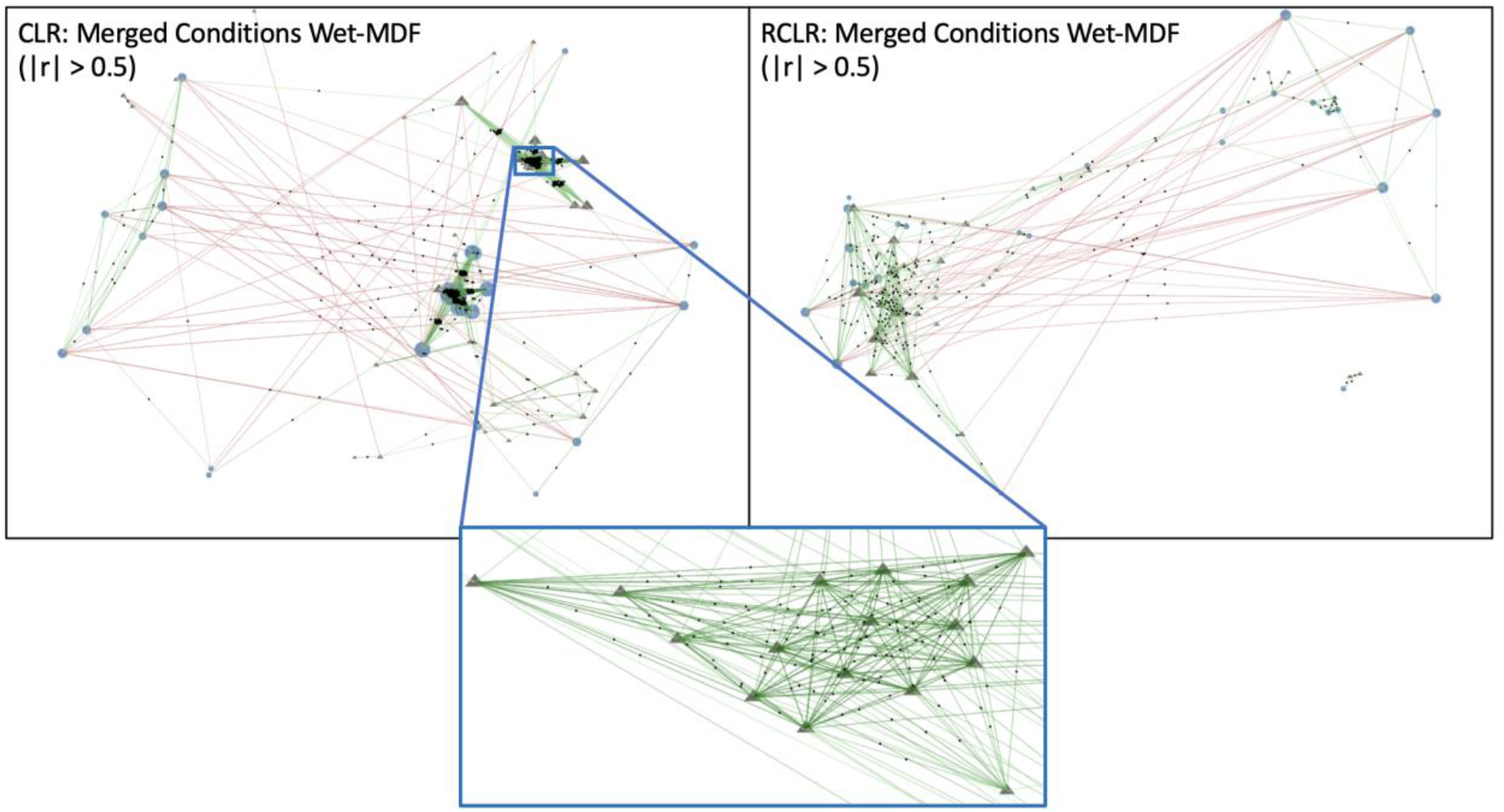
Results of bootstrap network analysis for the same datasets after clr (LEFT) and rclr (RIGHT) transformation. The dataset included bacterial and fungal genera counts based on 16S and ITS sequencing of microbial communities growing on the common building material medium density fiberboard (MDF) that was allow to grow at constant high humidity for several weeks. The circles indicate bacterial genera and the triangles indicate fungal genera. Green lines indicate positive correlations (Spearman r > 0.5) and red lines negative (Spearman rank r < 0.5) after 1000 bootstrap replicates using pybootnet (23). The box insert shows zooms in on a portion of the clr transformed network of highly artefactually correlated fungal genera (r-values between 0.75 and 1.0) that do not appear in the rclr transformed network.

Given our results, we strongly recommend using rclr transformation instead of clr transformation for correlation network analyses to avoid false correlations, especially highly sparse datasets. These correlations were strong even bootstrap analysis which typically helps reduce spurious correlations (23). Replacing zeros with any value prior to transformation will lead to the same results since the geometric mean of each sample will be different.

